# CD44/Integrin β1 association drives fast motility on HA substrates

**DOI:** 10.1101/2024.05.18.594819

**Authors:** Tanusri Roy, Sarbajeet Dutta, Lakshmi Kavitha Sthanam, Shamik Sen

## Abstract

In addition to proteins such as collagen (Col) and fibronectin, the extracellular matrix (ECM) is enriched with bulky proteoglycan molecules such as hyaluronic acid (HA). However, how ECM proteins and proteoglycans collectively regulate cellular processes has not been adequately explored. Here, we address this question by studying cytoskeletal and focal adhesion organization and dynamics on cells cultured on polyacrylamide hydrogels functionalized with Col, HA and a combination of Col and HA (Col/HA). We show that fastest migration on HA substrates is attributed to the presence of smaller and weaker focal adhesions. Integrin *β*1 co-localization and its association with CD44—which is the receptor for HA, and insensitivity of cell spreading to RGD on HA substrates suggests that focal adhesions on HA substrates are formed via integrin association with HA bound CD44. Consistent with this, adhesion formation and cell motility were inhibited when CD44 was knocked out. Collectively, our results suggest that association of integrin *β*1 with CD44 drives fast motility on HA substrates.

## Introduction

The extracellular matrix (ECM) of tissues represents the binding substrate for all adherent cells in the body^1^. Cells dynamically respond to the biomechanical properties of the ECM by modulating their cellular processes and cytoskeletal organization^2^. Cellular processes ranging from the division, migration to differentiation are regulated by the microenvironment which provides a combination of soluble and insoluble cues^3^. While the soluble components mostly comprise of signaling molecules, the insoluble components include fibrous proteins and glycosaminoglycans (GAGs)^4^. Adherent cells recruit integrin machinery to sense the elasticity and composition of the fibrous ECM network and upregulate downstream signaling networks for maintaining cellular homeostasis^5^. Mechanoadaptation represents the process by which cells alter their own biophysical properties in response to physical properties of the microenvironment^6^. In a seminal work, Janmey and co-workers demonstrated that when plated of substrates of stiffness less than 5 kPa, NIH 3T3 fibroblasts match their cortical stiffness to that their substrate^7^. Subsequently, several studies have illustrated this behavior in other cell types including that of cancer cells^8^. While the role of integrins in mediating mechanoadaptation is clear, cellular adaptation to GAGs is not well understood.

GAGs are long linear/branched polysaccharide chains which are abundantly present at cell-cell and cell-ECM junctions^9^. Most of the GAGs are sulphated polysaccharides such as chondroitin sulphate (CS), heparin/heparan sulphate (HS), and dermatan sulphate (DS). They are covalently attached to different ECM proteins making them the proteoglycans (PGs)^10^. Contrary to the abovementioned GAGs, Hyaluronic Acid (HA) is the only non sulphated GAG which is not part of any PG but is responsible for the aggregation of different PGs. PGs with unique sulphated patterns not only contribute to the structural heterogeneity of the tissue but also to the unique functions of GAGs^11^. While sulphated GAGs form proteoglycans, HA due its high molecular weight and viscosity is more than a joint lubricant of tissues^12^. Cluster determinant 44 (CD44) is the most important receptor ligand for HA along with other receptors like toll-like receptor 4 (TRL4)^13^. CD44 is expressed at the early stages of embryonic development in both mouse and human stem cells and binding of CD44 to HA activate multiple signaling pathways from pro survival signaling pathways, migration to homing of stem cells^14^.

Though the contribution of individual receptors of ECM proteins (i.e., integrins) and those of HA (i.e., CD44) have been well studied^11,12^, the collective effect of both these receptors in mediating adhesion, spreading and cytoskeletal organization in complex microenvironments has not been adequately explored. Recently, we showed that mouse embryonic stem cells which maintain pluripotency on collagen-coated substrates, are unable to adhere to HA gels alone, but adhere and differentiate robustly into neurogenic lineage on substrates functionalized with a combination of collagen (Col) and HA^17^. While this study demonstrates the effectiveness of these two ligands in inducing differentiation when presented together, the existence of any crosstalk between the two receptor systems (i.e., integrins and CD44), and its importance remains unclear. Since differentiation is closely associated with cell mechanics^13^, here we asked if interaction between CD44 and integrins can modulate cell mechanics. By studying spreading, motility and cytoskeletal organization of mouse embryonic fibroblasts (MEFs) and HT-1080 human fibrosarcoma cells on substrates coated with Col, HA and a combination of Col and HA in 1:1 ratio, we show that Col elicits a stronger mechanoadaptation response compared to HA, with both integrins and CD44 collectively mediating mechanoadaptation on (Col + HA) substrates. Furthermore, our results suggest that the integrin subunit – β1 associate with CD44 in ligand-dependent manner at adhesions for maintaining cell homeostasis.

## Results

### Stiffness-dependent cell spreading is muted on HA substrates

As a first step towards probing how extracellular matrix (ECM) stiffness regulates cell morphology, PA gels of varying stiffnesses were fabricated and functionalized with collagen (Col), hyaluronic acid (HA) and a combination of collagen and hyaluronic acid (Col/HA). Varying the ratio of acrylamide:bis-acrylamide allowed us to obtain gels of stiffness ranging from soft 0.6 kPa gels to moderately stiff 33 kPa gels (Fig. 1 Ai-iii). Fluorescence images and cryo-SEM images of functionalized PA gels confirmed successful coating of collagen and hyaluronic acid (HA) of the gels (Fig. 1 B and Supp. Fig.1 A), without any alteration of the gel stiffness (Fig. 1 C and Supp. Fig. 1 B). When cultured on the three ligand-coated PA gels of varying stiffnesses, mouse embryonic fibroblasts (MEFs) exhibited ligand-dependent spreading responses. MEFs displayed a high basal level spreading on Col-coated substrates which was comparable on 0.6 kPa and 4 kPa stiffnesses but further increased on the 33 kPa gels (Fig. 1 Di, Dii). However, cell spreading reduced significantly on both the HA and Col/HA coated gels in comparison to Col-coated gels, with weaker stiffness sensitivity. Together, these results indicate that stiffness-dependent spreading is ligand-dependent, with minimal spreading on HA-coated gels.

**Figure 1:**
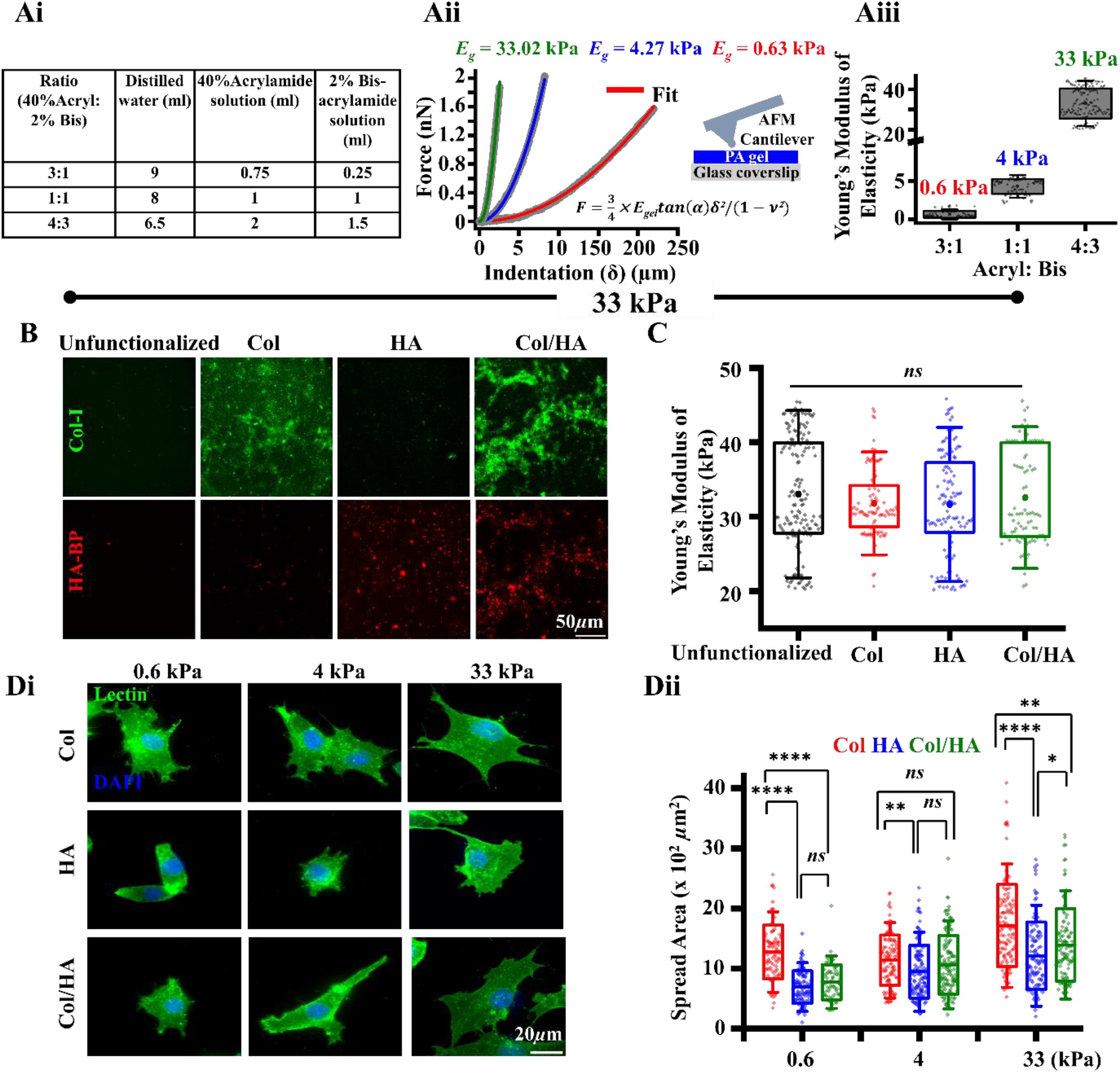
Fabrication, characterization of polyacrylamide (PA) hydrogels and mechanoadaptation of mouse embryonic fibroblasts (MEFs) on collagen (Col), hyaluronic acid (HA) and Col/HA coated PA gels: **(Ai)** Table showing different ratios of 40% acrylamide and 2% bis-acrylamide solution **(Aii)** Representative force-indentation curves of PA gels of varying acrylamide: bisacrylamide (acryl: bis) concentrations probed using Atomic fore microscopy (AFM). Raw force-indentation curves were fitted with Hertz model to obtain estimates of substrate stiffness. **(Aiii)** Quantification of PA gel stiffness (*n* ≥ 60 indentations per condition across *N* = 3 independent experiments) with varying acryl: bis concentrations. Error bars represent SD. **(B)** Representative immunofluorescence images of unfunctionalized and Col, HA and Col/HA functionalized PA gels of 4:3 acryl: bis stained with Col-I antibody and HA binding peptide (HA-BP). Scale Bar = 50 *µ*m. **(C)** Quantification of substrate stiffness of unfunctionalized, Col, HA and Col/HA functionalized PA gels (*n* ≥ 60 indentations per condition across *N* = 3 independent experiments). Error bars represent SD. **(Di)** Representative lectin-stained images of MEFs on Col, HA and Col/HA functionalized 0.6 kPa, 4 kPa and 33 kPa PA gels after 24 hrs of culture. Scale Bar = 20 *μ*m. **(Dii)** Quantification of cell spreading area on Col, HA and Col/HA functionalized 0.6 kPa, 4 kPa and 33 kPa PA gels after 24 hrs of culture. (*n* ≥ 30 cells per condition across *N* = 3 independent experiments). Error bars represent SD. One-way Anova with Tukey’s test was used for comparing means (\**p*-value ≤0.05, ** *p*-value ≤0.01, **** *p*-value ≤0.0001, ns = non-significant *p*-value>0.05).

### Weaker adhesions enable faster migration on HA substrates

Since highest spreading for all the three ligands was observed on 33 kPa gels, subsequent experiments were done on these gels only. To check the effect of ligand composition on cell motility, cells were tracked for 12 hrs under the microscope. Surprisingly, analysis of single cell trajectories revealed highest motility on HA substrates and comparable motility profiles on Col and Col/HA substrates (Fig. 2 Ai, ii, Supp Movie 1). To probe the role of altered tensional homeostasis in regulating cell spreading and motility, cells were stained with phalloidin to mark the F-actin stress fibers. While cells on HA gels formed shorter stress fibers in comparison to the other conditions (Fig. 2 Bi, ii), immunostaining and western blotting of active contractility using pMLC revealed increased pMLC on HA substrates, with highest levels observed on the Col/HA gels (Fig. 2 Ci, ii).

**Figure 2:**
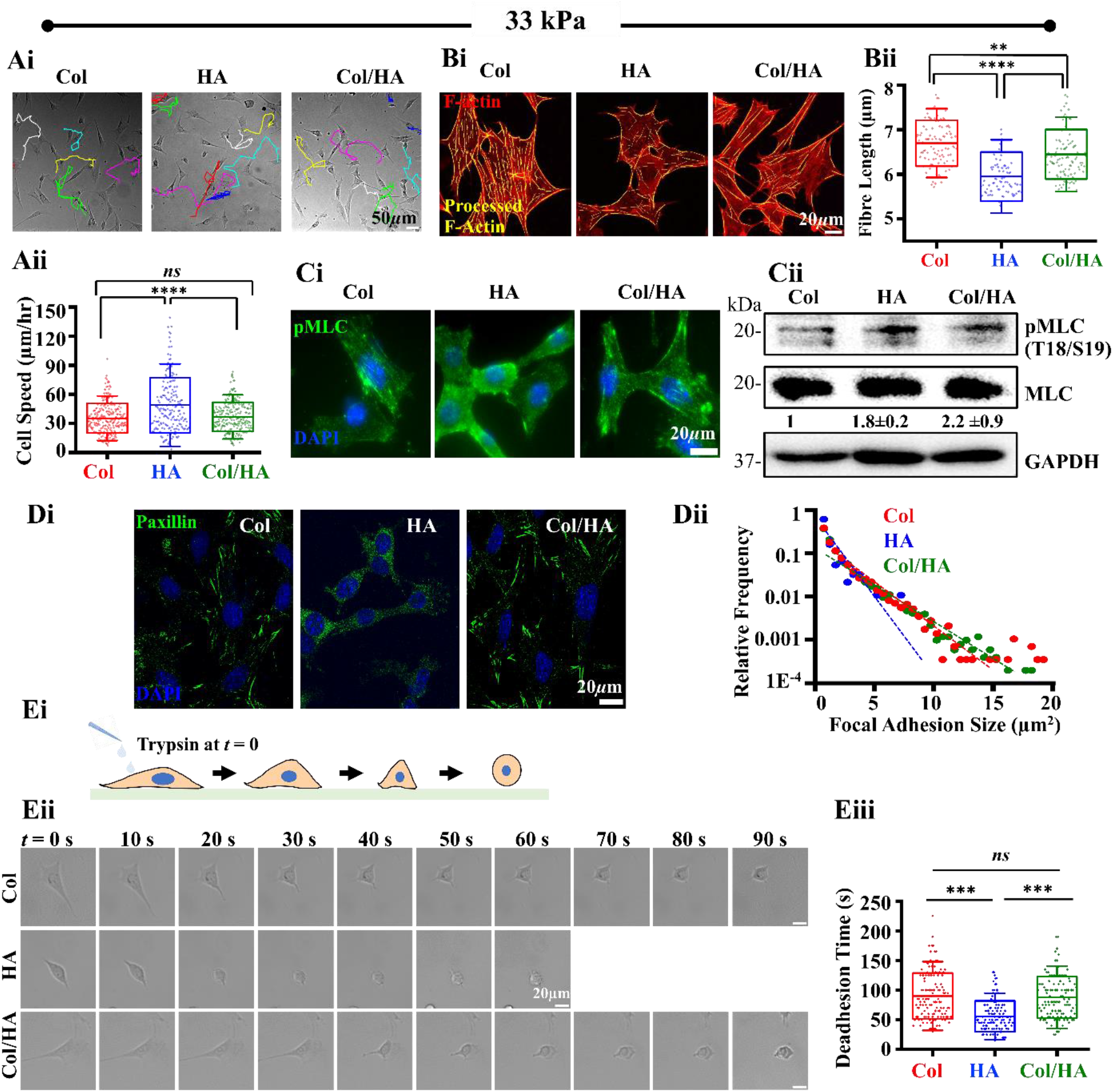
Migration, cytoskeletal organization and adhesion of MEFs on Col, HA and Col/HA functionalized 33 kPa PA gels: **(Ai)** Representative random cell migration trajectories of individual cells on Col, HA and Col/HA functionalized PA gels. Scale Bar = 50 *μm*. **(Aii)** Quantitative analysis of cell speed on Col, HA and Col/HA functionalized PA gels (*n* ≥ 30 cells per condition across *N* = 3 independent experiments). Error bars represent SD. **(Bi)** Representative processed images of phalloidin stained F-actin images of MEFs cultured on Col, HA and Col/HA functionalized PA gels. Scale Bar = 20 *µ*m **(Bii)** Quantification of F-actin fibre length per cell (*n* ≥ 30 cells per condition across *N* = 3 independent experiments). Error bars represent SD. **(Ci)** Representative immunofluorescence images of pMLC2(S19) on MEFs cultured on Col, HA and Col/HA functionalized PA gels. Scale Bar = 20 µm. **(Cii)** Western blots and quantitative analysis ±SEM of pMLC2 (T18/S19) /MLC on MEFs cultured on Col, HA and Col/HA functionalized PA gels (*N* = 3 independent experiments). **(Di)** Representative immunofluorescence images of paxillin staining in MEFs cultured on Col, HA and Col/HA functionalized PA gels. Scale Bar = 20 *µ*m. **(Dii)** Size distribution of paxillin adhesions in MEFs cultured on Col, HA and Col/HA functionalized PA gels (*n* ≥ 30 cells per condition across *N* = 3 independent experiments). **(Ei)** Schematic of trypsin de-adhesion. **(Eii)** Representative phase-contrast images of MEFs upon addition of trypsin at time *t* = 0. Scale Bar = 20 *µ*m. **(Eiii)** Quantification of de-adhesion time (*n* ≥ 30 cells per condition across *N* = 3 independent experiments). Error bars represent SD. One-way Anova with Tukey’s test was used for comparing means (\**p*-value ≤0.05, *** *p*-value ≤0.001, **** *p*-value ≤0.0001, ns = non-significant *p*-value>0.05).

Since cell motility is dependent on both focal adhesion formation and their turnover, we probed how focal adhesions were altered across the three conditions. Though paxillin-positive focal adhesions were detected on all the three substrates, analysis of focal adhesion size distribution revealed the formation of smaller focal adhesions (<10 *μm*^2^) on HA substrates; in comparison, cells cultured on Col and Col/HA gels possessed larger focal adhesions (upto 20 *µm*^2^) (Fig. 2 Di, ii), suggesting that faster adhesion turnover may mediate faster migration on HA substrates. To test this, we made used of trypsin de-adhesion assay wherein adherent cells are incubated with warm trypsin and their detachment kinetics tracked (Fig. 2 Ei, Supp Movie 2) ^14,15^. Faster de-adhesion may be associated with weaker adhesions and higher cell contractility. In line with formation of smaller number and size of adhesions, cells de-adhered fastest on the HA substrates (Fig. 2 Eii, iii). Collectively, these results suggest formation of smaller and weaker adhesions combined with elevated levels of actomyosin contractility drive faster migration on HA substrates.

### Focal adhesions on HA substrates may be mediated via integrin-CD44 interactions

What might be the nature of focal adhesions on HA substrates? Unlike Col and Col/HA gels where integrins can directly bind to collagen on these two substrates, adhesion on HA is likely to be mediated by its receptor, the transmembrane glycoprotein CD44^16–18^. Since integrin β1 is known to bind to CD44, we speculate focal adhesion formation on HA substrates is mediated by integrin associated with HA engaged CD44. Furthermore, this indirect coupling may be the reason for weaker adhesion formation on HA substrates. To test our hypothesis, integrin-β1 (ITGβ1) and CD44 whole cell expression levels were probed on Col, HA and Col/HA-coated 33 kPa PA gels. Intriguingly, comparable levels of both ITGβ1 and CD44 were observed from whole cell lysates across all the three substrates (Fig. 3A). To probe if integrin signaling is activated on these substrates, pFAK(Y397) levels were probed by immunostaining and quantified by western blotting of whole cell lysates. Interestingly, pFAK/FAK quantification showed increased activation on HA substrates compared to Col-coated substrates (Fig.3 Bi, Bii), suggesting that integrin signaling on HA substrates could be activated by CD44-integrin interaction on the HA substrates. To test integrin-CD44 interactions on these substrates, cells were immunostained with both ITGβ1 and CD44 wherein merged images displayed colocalization across all three substrates (Fig. 3 C), presumably indicating the role of CD44 in mediating cell migration on HA substrates through integrin association. To test the role of CD44 in cell spreading via integrins, RGD peptide was added to cells to block integrin binding to its substrates while RGE served as control (Fig. 3 Di). Integrin blocking for 2 hrs severely reduced cell spreading on both Col and Col/HA substrates but did not affect cell spreading on HA substrates (Fig. 3 Dii, iii). Thus, cell spreading and migration on HA substrates is mediated through CD44 engaging via integrin signaling. Collectively, these results suggest that CD44 could possibly interact with integrin β1 to activate integrin signaling forming “weak” focal adhesions with higher turnover on HA substrates that triggers faster migration in comparison to Col or Col/HA substrates.

**Figure 3:**
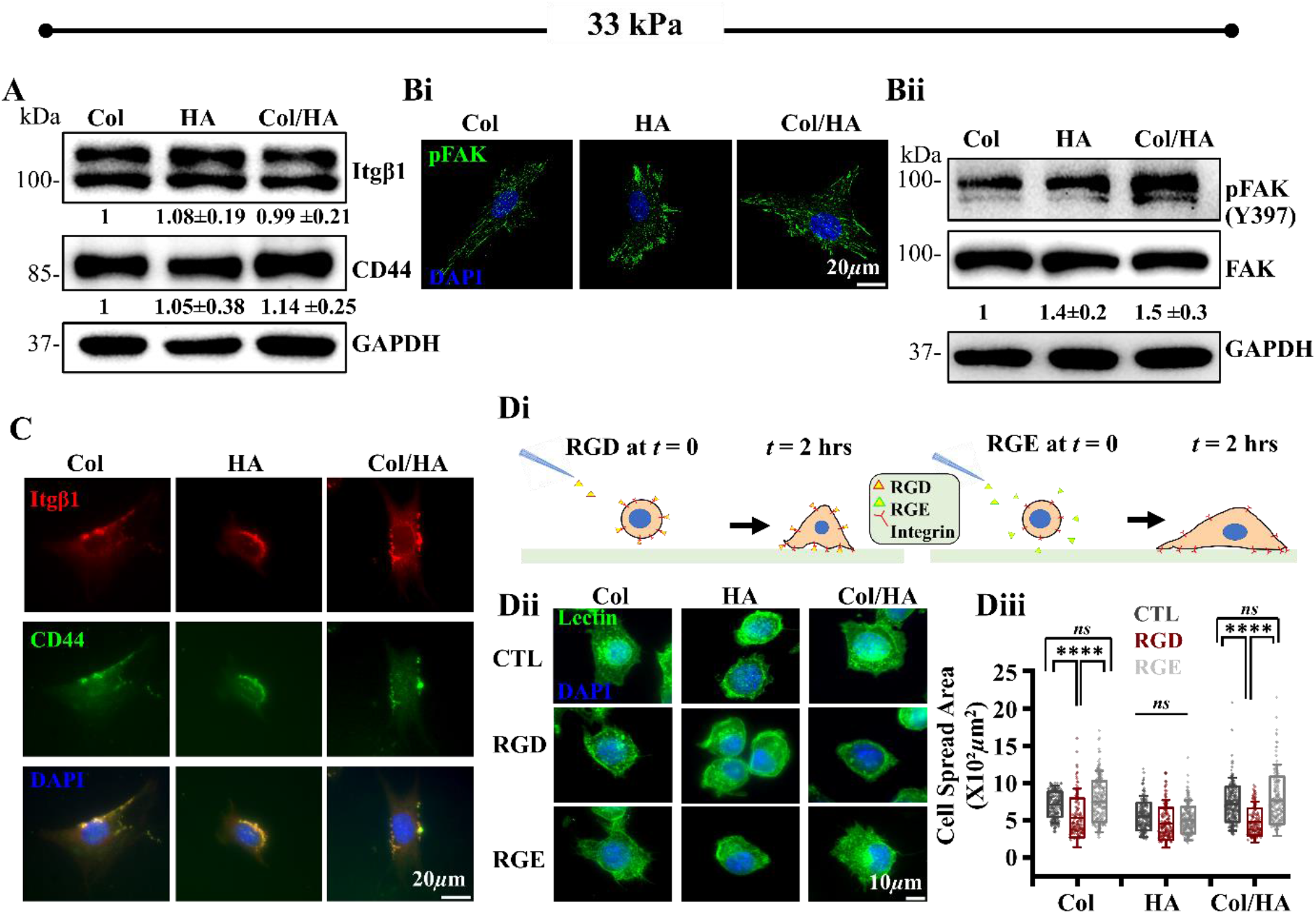
CD44 expression and localization at focal adhesions: **(A)** Western blots and quantitative analysis (±SEM) of ITGβ1 and CD44 expression on MEFs cultured on Col, HA and Col/HA functionalized PA gels (*N* = 3 independent experiments). **(Bi)** Representative immunofluorescence images of pFAK (Y397) on MEFs cultured on Col, HA and Col/HA functionalized PA gels. Scale Bar = 20 *µ*m. **(Bii)** Western blots and quantitative analysis (±SEM) of pFAK/FAK (Y397) on MEFs cultured on Col, HA and Col/HA functionalized PA gels (*N* = 3 independent experiments). **(C)** Representative immunofluorescence images of ITGβ1(red) and CD44 (green) on MEFs cultured on Col, HA and Col/HA functionalized PA gels. Scale Bar = 20 *µ*m. **(Di)** Schematic representation of RGD/RGE treatment on MEFs cultured on Col, HA and Col/HA functionalized PA gels. **(Dii)** Representative images of untreated and RGD/RGE treated or untreated MEFs, fixed at t = 2 hrs and stained with lectin. Scale Bar = 10 *µ*m. **(Diii)** Quantification of MEF spread area on Col, HA and Col/HA functionalized PA gels after 2 hrs of RGD/RGE treatment (*n* ≥ 30 cells per condition across *N* = 3 independent experiments). Error bars represent SD. One-way Anova with Tukey’s test was used for comparing means (***** *p*-value ≤0.0001, ns = non-significant *p*-value>0.05). For all blots, GAPDH serves as loading control.

### CD44/Integrin β1 association drives fast migration on HA substrates

Thus far, our results of cell spreading on the different ligand-coated substrates are indicative of more robust spreading on Col-coated substrates compared to that on HA-coated substrates. However, faster migration due to weak focal adhesions observed on HA-coated substrates raises the possibility that CD44-based adhesions determine cell migration through integrin association. Since MEFs form small adhesions on HA-coated substrates, for ease of experiments, we performed experiments with HT1080 fibrosarcoma cells which are much larger than MEFs and also express higher levels of Itgβ1 and CD44 (Fig. 4A). Co-immunostaining of Itgβ1 and CD44 on Col, HA and Col/HA gels revealed Itgβ1/CD44 co-localization across all three gels (Fig.4 Bi, ii). To check for possible association between Itgβ1 and CD44, we performed proximity ligation assay (PLA) (Fig. 4Ci). In contrast to negative controls, i.e., IgG and 2° only, prominent PLA signal was detected on Col, HA and Col/HA substrates (Fig. 4Cii). However, quantification of PLA intensities revealed strongly Itgβ1/CD44 association on HA gels (Fig. 4Ciii). Collectively, these results establish robust association between Itgβ1 and CD44 on HA coated gels.

**Figure 4:**
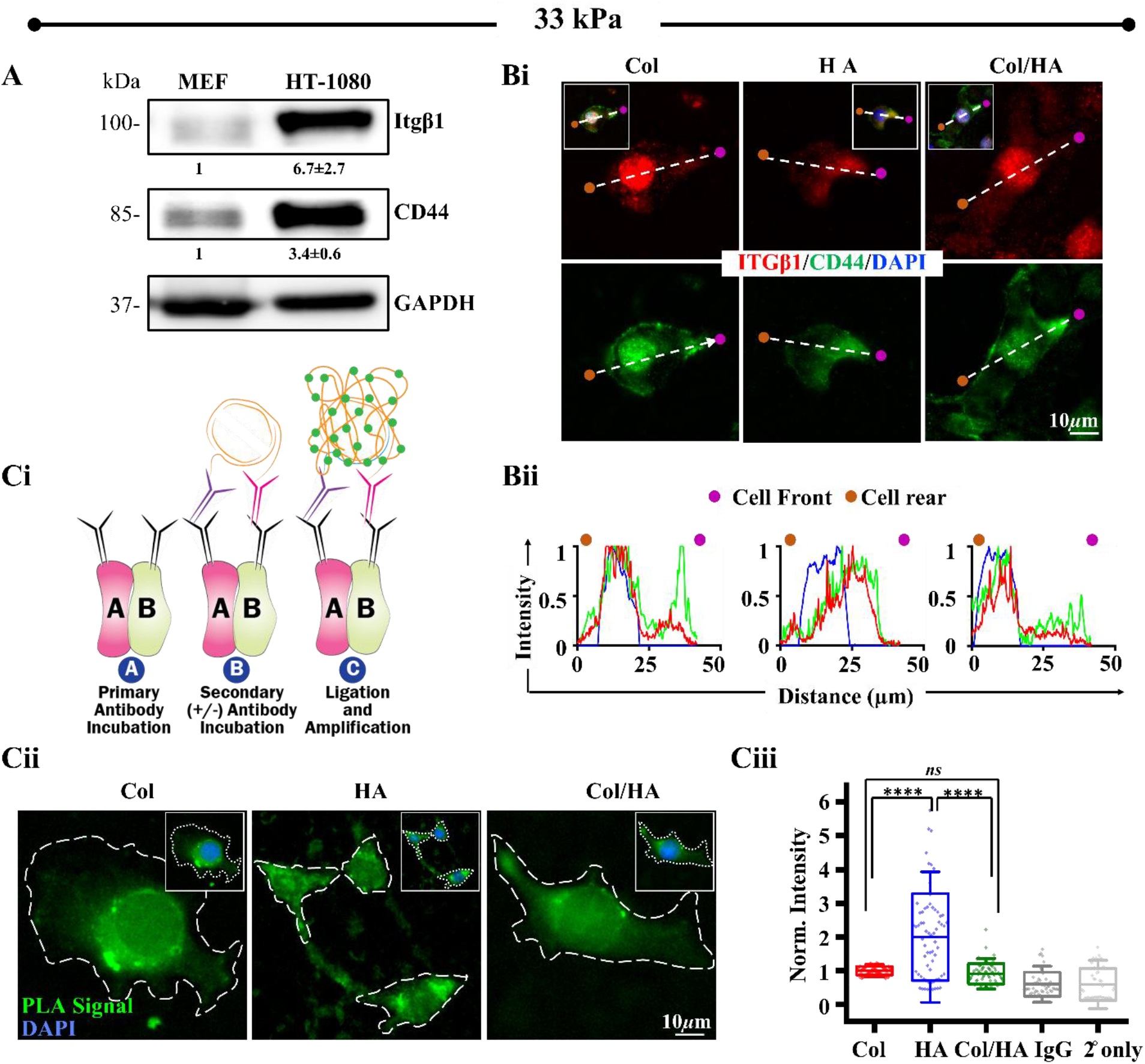
Interaction between ITGβ1 and CD44: **(A)** Western blots and quantitative analysis (±SEM) of ITGβ1 and CD44 on MEFs and HT-1080 human fibr0sarcoma cell lines (*N* = 3 independent experiments). GAPDH serves as loading control. **(Bi)** Representative immunofluorescence images of ITGβ1(red) and CD44 (green) in HT-1080 cells on Col, HA and Col/HA functionalized PA gels. Scale Bar = 10 *µ*m. **(Bii)** Representative intensity plots of ITGβ1, CD44 and DAPI depicting their co-localization pattern. **(Ci)** Schematic representation of proximity ligation assay (PLA). **(Cii)** Representative PLA signals depicting ITGβ1/CD44 interaction in cells cultured on Col, HA and Col/HA functionalized substrates. Scale Bar = 10 *µ*m. **(Ciii)** Quantification of PLA intensities of cells cultured on Col, HA and Col/HA functionalized PA gels along with signals on IgG and secondary only (2° only) controls. Intensities were normalized to that of cells on Col coated substrates (*n* ≥ 20 cells per condition across *N* = 3 independent experiments). Error bars represent SD. One-way Anova with Tukey’s test was used for comparing means (***** *p*-value ≤0.0001, ns = non-significant *p*-value>0.05).

To test the functional importance of CD44-Itgβ1 association on HA substrates, KO-CD44 HT-1080 cells were established using Crispr-Cas9, and validated by western blotting of whole cell lysate of WT and CD44-KO cells (Fig. 5A). Though CD44 was dispensible for cell spreading on Col and Col/HA gels, surprisingly, CD44-KO cells were found to spread more than WT cells (Fig. 5 Bi, ii). While CD44-KO did not impede cell motility on Col and Col-HA gels, a significant drop in motility was observed on HA substrates (Fig. 5Ci, ii, Supp Movie 3). Quantification of focal adhesion size revealed robust reduction in the number of focal adhesions formed on HA substrates in CD44-KO cells, and also on Col/HA gels (Fig. 5Ci, ii). Further, quantification of focal adhesion turnover on glass coverslips revealed drop in adhesion stability on both HA and Col/HA substrates in CD44-KO cells (Fig. 5Di, ii, Supp Movie 4). Collectively, these results highlight the importance of CD44-Itgβ1 association in sustaining cell migration on HA substrates via regulation of adhesion size and dynamics.

**Figure 5:**
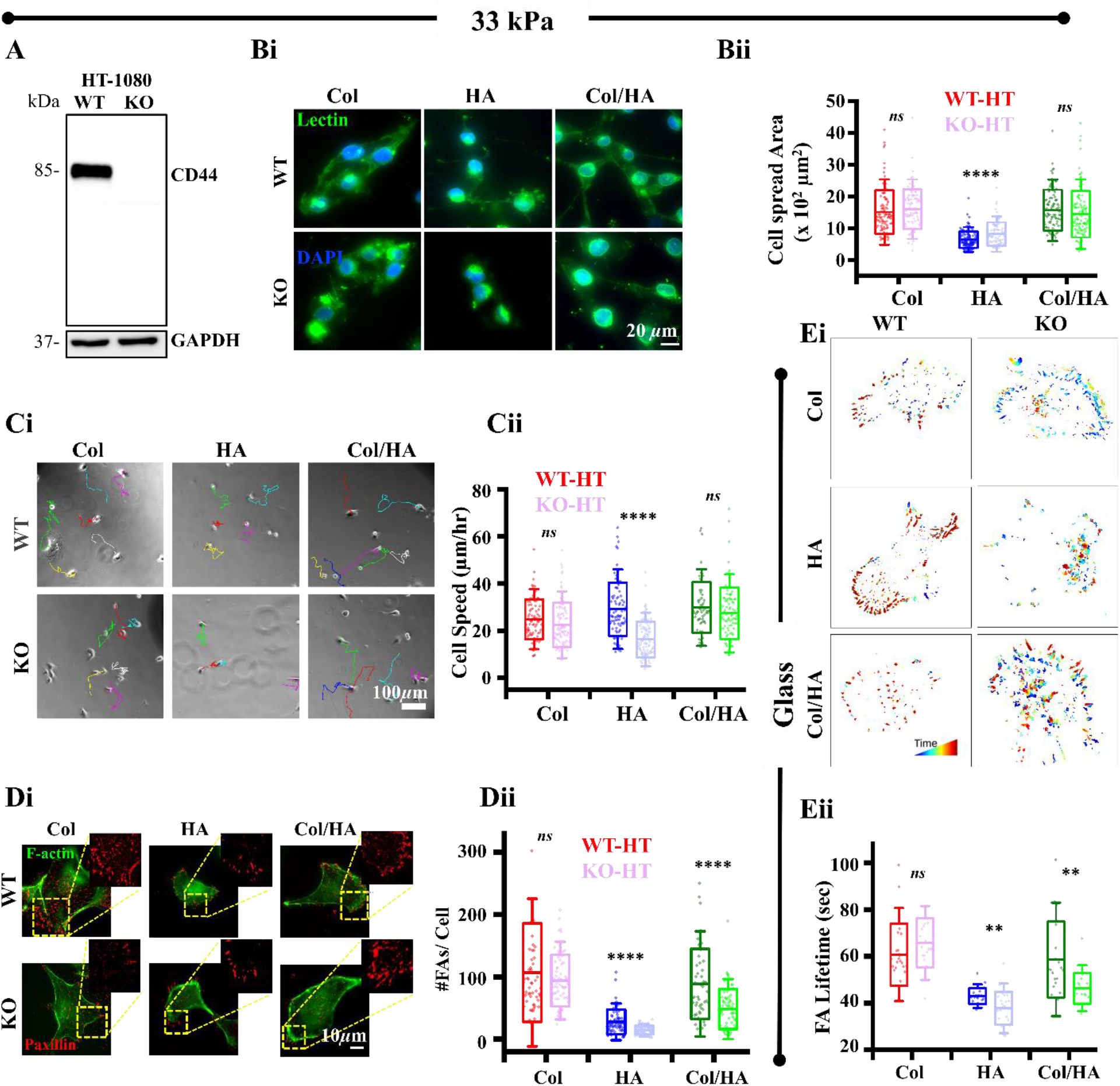
Cell spreading, migration and adhesion of CD44-KO cells Col, HA and Col/HA-coated substrates: **(A)** Validation of CD44-KO by western blotting of WT vs CD44-KO HT-1080 cells. GAPDH serves as loading control. **(Bi)** Representative lectin-stained images of WT vs CD44-KO cells on Col, HA and Col/HA functionalized 33 kPa PA gels. Scale Bar = 20 *µ*m. **(Bii)** Quantification of cell spread area of WT vs CD44-KO cells on Col, HA and Col/HA functionalized 33 kPa PA gels. (*n* ≥ 30 cells per condition across *N* = 3 independent experiments). Error bars represent SD. One-way Anova with Tukey’s test was used for comparing means. **(Ci)** Representative random cell migration trajectories of individual WT vs CD44-KO cells on Col, HA and Col/HA functionalized PA gels. Scale Bar = 100 *μ*m. **(Cii)** Quantitative analysis of WT vs CD44-KO cell speed on Col, HA and Col/HA functionalized PA gels (*n* ≥ 30 cells per condition across *N* = 3 independent experiments). Error bars represent SD. One-way Anova with Tukey’s test was used for comparing means. **(Di)** Representative F-actin (green) and paxillin (red) co-stained images of WT vs CD44-KO cells on Col, HA and Col/HA functionalized PA gels. Scale Bar = 10 *µ*m. **(Dii)** Quantification of focal adhesion count per cell (paxillin) of WT vs CD44-KO cells on Col, HA and Col/HA functionalized PA gels. (*n* ≥ 30 cells per condition across *N* = 3 independent experiments). Error bars represent SD. One-way Anova with Tukey’s test was used for comparing means. **(Ei)** Representative focal adhesion turnover plots of WT vs CD44-KO cells transfected with eGFP-paxillin on Col, HA and Col/HA coated glass. **(Eii)** Quantification of focal adhesion turnover of WT vs CD44-KO cells on Col, HA and Col/HA coated glass. (*n* ≥ 5) cells per condition across *N* = 3 independent experiments). Error bars represent SD. One-way Anova with Tukey’s test was used for comparing means (**** *p*-value ≤0.0001, \*\**p*-value ≤0.01, ns = non-significant *p*-value>0.05).

## Discussion

This study illustrates the importance of the cell surface glycoprotein receptor CD44 in driving fast migration on HA substrates through its association with Itgβ1 (Fig. 6). Reduced spreading and weaker adhesions on HA substrates as compared to Col and Col/HA substrates suggests that weak adhesions may enables cells to migrate faster on HA substrates. Although Itgβ1 and CD44 levels remain unchanged across these ligands, faster migration on HA substrates correlates with activation of downstream integrin signaling as observed from pFAK images and blots. Using PLA, we demonstrate the association of Itgβ1 with CD44, prominently on the HA coated gels. Loss of CD44 significantly reduced cell motility by lowering focal adhesion number and turnover, underlining the importance of CD44-Itgβ1 association in sustaining fast migration of cells on HA coated substrates.

**Figure 6:**
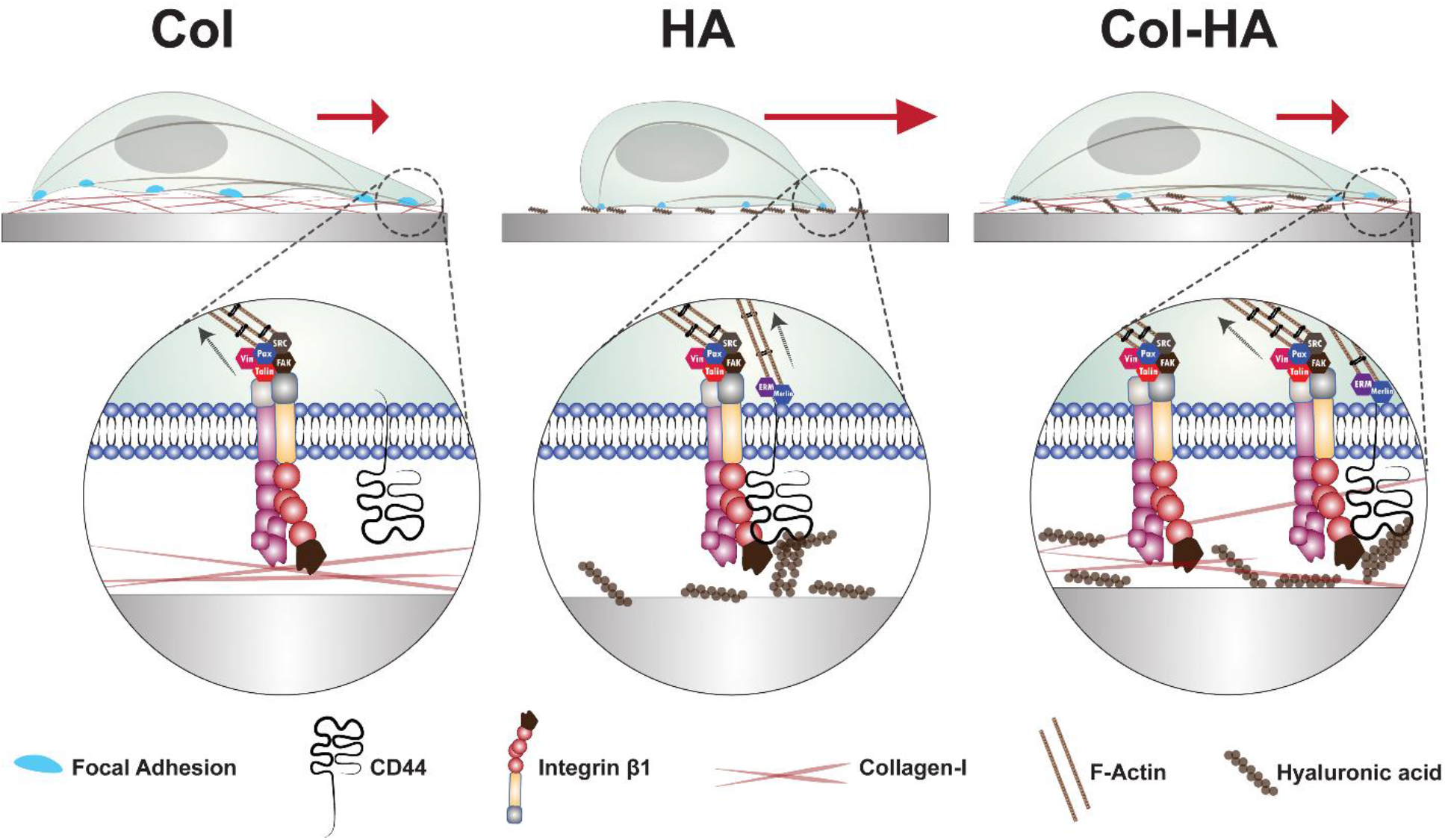
Schematic summary representing CD44-Integrin β1 association leading to faster migration on HA coated gels.

HA is secreted by cells of mesenchymal origin into the ECM that performs various physiological functions from cell adhesion to differentiation ^19,20^. HA binding to CD44 triggers conformational changes allowing its intracellular domain binding to cytoskeletal elements, and activation of downstream signalling pathways^16,17^. Unlike synthetic polymers, HA is biologically active and its water retaining capacity enables maintenance of viscoelastic properties of connective tissues ^20–22^. In addition, soft HA gels crosslinked with either Col or laminin showed similar or higher extent of spreading and mechanoadaptation comparable to that of stiff PA gels coated with Col or laminin^23^ suggesting that the cell mechanosensitivity on elastic substrates is greatly dependent on the ligand presented rather than the stiffness of the substrate alone^24^. This suggests that CD44-based adhesions which are not primary adhesion receptors have the ability to alter the mechanoadaptation of cells when cultured on HA substrates alone. Consistent with these observations, cells cultured on HA substrates are relatively less spread when compared to those on Col substrates.

CD44 is an accepted marker for breast cancer cells, HA being its primary ligand ^25,26^. The role of CD44 in regulation of integrin expression and mediating cell-matrix adhesion has been reported in literature^27^. Human mesenchymal stem cells (hMSCs) have been documented to migrate to liver injury sites in Itgβ1-CD44 dependent manner^28^. Apart from these observations, the distribution of high to low molecular weight HA in various tissues, and differences in the expression of CD44 and recycling of CD44 in response to the high/low molecular HA raises a very interesting phenomenon of structural complexity regulating cellular adhesion and mechanics^18^. Consistent with our observations, HA have been reported to promote CD44 mediated activation of talin and paxillin through Itgβ1α5 receptor in breast cancer cells^27^. Furthermore, Itgβ3 has been shown to play a key role in CD44 mediated activation of NFκB pathway during lung cancer cell growth^29^.

Even though integrins co-localize with CD44 at adhesions, due to absence of integrin-binding motifs on HA, mechanoadaptation is mediated exclusively by CD44. However, the strength of this mechanoadaptation is weaker than that mediated by direct integrin-matrix adhesions, as evidenced by lesser extent of spreading, and faster de-adhesion on HA substrates. This is also clear from the presence of shorter stress fibers and lesser focal adhesions in cells cultured on HA substrates in comparison to the extensive stress fiber network and larger adhesions observed on Col substrates. However, prominent activation of FAK illustrates that even membrane stabilization of Itgβ1 by CD44 is sufficient to activate downstream adhesion signaling. While reduced spreading on Col and Col/HA gels in the presence of RGD is consistent with the integrin blocking activity of RGD via competitive binding, insensitivity of cell spreading to RGD on HA substrates suggests that even RGD bound Itgβ1 can remain CD44 associated and sustain adhesion signaling.

In conclusion, in addition to demonstrating the collective influence of Col and HA in regulating cell mechanics, our observation of association of CD44 with integrin β1 in a ligand-dependent manner illustrates the active crosstalk between CD44 and integrins that determines cell migration substrates devoid of collagen as ECM. Future studies aimed at characterizing the nature and mechanisms of crosstalk between these two receptors can further improve our understanding of how cells regulate tensional homeostasis by modulating the expression, localization and interaction between these two receptors.

## Materials & Methods

### Cell culture & Reagents

Mouse embryonic fibroblasts (MEFs), HT-1080 human fibrosarcoma cells and CD44-KO HT-1080 cells were cultured in DMEM (with phenol red) supplemented with 10% FBS (Invitrogen) and 1X antibiotic-antimycotic at 37°C and 5% CO_2_ concentration. Media was changed once every two days after washing with PBS (Sigma). For experiments, cells were dislodged with 1x trypsin (HiMedia) and cultured on polyacrylamide (PA) hydrogels coated with rat tail Collagen type I (Sigma) and/or low molecular weight (8-15 kDa) hyaluronic acid (HA, SRL). For integrin blocking experiments, cells were incubated with either RGD or RGE (SRL) at a concentration of 2 mg/ml for 2 hours before imaging.

### CD44-KO cell line generation

CRISPR CAS9 mediated knockout of CD44 was performed in HT1080 cells by designing sgRNAs against CD44 exon using the online available software (https://chopchop.cbu.uib.no/) and cloning into pSpCas9(BB)-2A-Puro (PX459) V2.0 plasmid (Addgene Plasmid# 62988, gift from Feng Zhang) according to the protocol described elsewhere^30^. Positively transfected HT1080 cells were selected by 1µg/mL puromycin treatment. The clonal propagation of the transfected cells was done by seeding one cell per well of a 96-well plate. The confirmation of the knockout of the propagated cells was determined by western blotting.

### Fabrication of polyacrylamide hydrogels

Polyacrylamide (PA) gels were fabricated by mixing acrylamide and bisacrylamide (BioRad) in varying ratios so as to obtain gels of stiffnesses of 0.6, 4.0 and 33 kPa as described previously elsewhere^31,32^. Gels were incubated with Sulpho-SANPAH (0.1mM, Pierce) dissolved in 50mM HEPES buffer, followed by UV light at 356nm. After washing with PBS, gels were incubated with rat tail Collagen type I at a concentration of 10µg/cm^2^ in PBS buffer. Overnight incubation was done for achieving uniform coating. For substrates coated with both collagen and HA, collagen and HA were combined at 1:1 ratio as described previously elsewhere^33^.

### Microscopy

cells were cultured on PA gels cast on 12mm coverslips at a seeding density of 7 x 10^3^ cells per well for 12 hours. For analysis of cell motility experiments, time lapse imaging was performed on a spinning disk confocal microscope (Zeiss) at 10x objective for 12 hours at 10 min intervals. Raw images were analyzed in Fiji ImageJ software quantification of cell motility.

For immunostaining of PA gels, the substrates were blocked using 5% BSA for 1 hour in room temperature (RT) following Col/HA coating. The gels were then incubated with anti-Col-I antibody and HA binding peptide (HABP) tagged with avidin overnight at 4° C. The following day, these gels were treated with alexa-fluor tagged secondary antibodies and streptavidin (Invitrogen) and then imaged under a fluorescent inverted microscope (Olympus, IX83).

For immunostaining of cells, they were fixed after 24 hours of culture using 4% PFA in PBS for 15 min, washed, blocked with 5% FBS for 1 hour in RT and incubated with one of the following antibodies overnight at 4 degrees: anti-pMLC (CST), anti-paxillin (Invitrogen), anti-pFAK (Invitrogen), anti-integrin β1 (Invitrogen) and anti-CD44 (Invitrogen). The following day, cells were incubated with alexa-fluor tagged secondary antibodies (Invitrogen) at RT for 2 hours and the nuclei were labelled using DAP (Sigma). Cells were stained with alexa-fluor tagged Lectin (Invitrogen) for analysis of cell spreading and with phalloidin (Invitrogen) for analysis of F-actin. Cells were imaged at 63x magnification using Scanning Probe Confocal Microscope (Zeiss, LSM 780) and 60 X magnification of Olympus IX83. Focal adhesion size and distribution was quantified according to the well-established protocol described elsewhere^34^. The cytoskeletal organization was quantified based on F-actin images using Filament Sensor plugin to determine the average length of actin stress fibers per cell per culture condition^35^.

### Trypsin de-adhesion assay

Cells cultured for 24 hours on the PA gels were washed with PBS followed by incubation with warm trypsin (Himedia). Images were acquired every 10 sec at 10x magnification until cells became rounded but were still attached to the underlying substrate as described earlier elsewhere ^14,36^. Deadhesion time (t) was measured until cells were rounded.

### AFM

Cell stiffness was measured using an Atomic Force Microscope (MFP3D, Asylum). Cells were probed slightly off-center with 10 kHz soft, pyramidal silicon nitride probes (Olympus) of nominal stiffness 20 pN/nm. The exact cantilever stiffness was determined using the thermal calibration method. Estimates of cell stiffness were obtained by fitting the first 500 nm of experimental force-indentation curves with Hertz model^37^. Similarly, the stiffness of gels was estimated by fitting the first 1000 nm of the experimental force-indentation curves as described elsewhere^32^.

### Proximity Ligation Assay (PLA)

Proximity Ligation Assay (PLA) was performed to detect the interactions between ITGβ1 and CD44 using Duolink® In Situ Red Starter Kit Mouse/Rabbit (DUO92101, Sigma-Aldrich) according to the manufacturer’s instructions. Briefly, The cells were fixed with 4% PFA for 10 min, washed twice with PBS, then cells were permeabilized with 0.25% Triton-X100 solution for 15 min followed by two times washing with PBS. The blocking solution supplied with the kit was used to block the cells for 1hr in a humid 37°C chamber. Then, anti-integrin β1 and anti-CD44 primary antibody incubation was done for overnight at 4°C. Next the cells were washed with 1x washing solution A twice. After that, Duolink PLUS/MINUS secondary antibody incubation was done in a humid 37°C chamber for 1hr. After two times washing with solution A, a ligation reaction was performed using the Duolink ligation regent for 30 min in a humid 37°C chamber followed by amplification with Duolink amplification regents for 2hr in the dark in a humid 37°C chamber. Final washing was performed using wash solution B and mounting was done using Duolink DAPI containing mounting media.

### Statistical Analysis

For statistical analysis, the normality of the data was first checked using the Kolmogorov-Smirnov normality test. Based on these results, either parametric or nonparametric tests were subsequently performed. Statistical significance was assessed using one-way ANOVA tests for parametric data, followed by Tukey’s posthoc test to compare the means. p < 0.05 being considered as statistically significant.

## Author contributions

Conceptualization: T.R., S.D., L.K.S., S.S.; Methodology: T.R., S.D., S.S.; Software: T.R., S.D., S.S; Validation: S.S.; Formal analysis: T.R., S.D., S.S.; Investigation: T.R., S.D., S.S.; Resources: S.S.; Data curation: T.R., S.D., S.S.; Writing - original draft: T.R., S.S.; Writing - review & editing: T.R., S.D., L.K.S., S.S.; Visualization: T.R., S.D., S.S.; Supervision: S.S.; Project administration: S.S.; Funding acquisition: S.S.

## Acknowledgements

This research had funding support from the Department of Science and Technology, Ministry of Science and Technology, India (DST/SJF/LSA-01/2016-17) and intramural funds provided by IIT Bombay. TR was supported by a CSIR fellowship (Grant # 09/087(0927)/2017-EMR-I) (Govt. of India) and SD was supported by a DBT fellowship (Grant # DBT/2017/IIT-B/850) (Govt. of India). Authors would also like to thank IRCC, IIT Bombay for providing Bio-AFM, confocal microscopy and Cryo-SEM facilities.

