## Supplementary Data for "CD44/Integrin β1 association drives fast motility on HA substrates"

Supplementary File

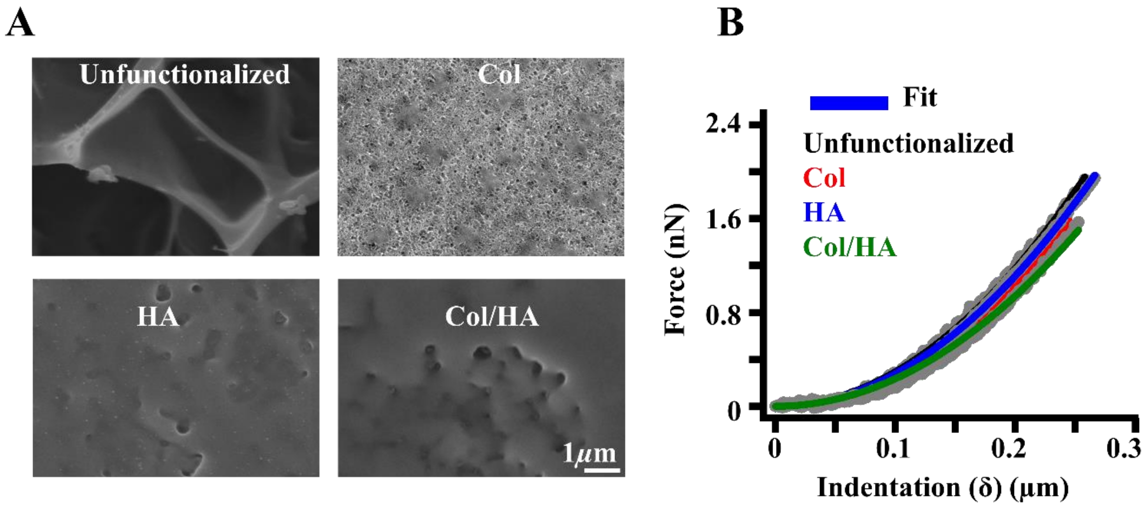


**Supplementary Figure 1: Col, HA and Col/HA coating on 33 kPa PA gels: (A)** Cryo-SEM images of uncoated and Col, HA, Col/HA coated PA gels visualized at 25,000X magnification. Scale Bar = 1$\mu$m. **(B)** Representative raw force-indentation curves fitted with Hertz model to estimate PA gel stiffness using AFM.

**Supplementary Videos:**

Supplementary Movie 1.mp4

Supplementary Movie 2.mp4

Supplementary Movie 3.mp4

Supplementary Movie 4.mp4
